# Targeted Searches for Novel Peptides in Big Mass Spectrometry Data Sets

**DOI:** 10.1101/239863

**Authors:** Yu Gao, Jiao Ma, Alan Saghatelian, John R. Yates

## Abstract

We present Post-Acquisition Targeted Searches (PATS), an easy-to-use tool that allows the identification of novel peptide/protein sequences from existing big mass spectrometry data sets. PATS filters out the unrelated peptidome before the time-consuming database search to significantly speed up the identification. Using interactome data sets, PATS visualizes protein interaction network and helps to assign putative functions to the target protein based on the “guilt by association” concept.

---

Protein identification generally consists of matching tandem mass spectrometry data to sequences in a database derived from DNA sequencing of an organism’s genome^1^. The protein database is dependent on our ability to predict open reading frames (ORFs) and, as a result, could be incomplete. Indeed, new techniques such as ribosome profiling and proteogenomics have revealed the existence of additional ORFs that are overlooked by traditional ORF-finding algorithms^2^. The existence of these non-annotated ORFs creates a new challenge to understand the biological functions of these proteins^3–5^.

Theoretically, large-scale mass spectrometry data from proteomic studies could be used to help assess the functions of additional proteins through data mining. For example, two maps of the human proteome^6, 7^ and several large-scale protein-protein interactome studies^8–10^ are the types of big data sets that could be used for data mining. However, the mass spectrometry data collected in these types of studies is often not exhaustively analyzed to identify all possible peptide sequences or modified peptides, and thus they constitute a potential resource to identify new information or to verify sequences obtained in other kinds of experiments. To mine these data sets, we must be able to perform searches with the newly discovered ORFs, peptides, or proteins.

The sheer size of the data and the huge computational complexity of these data sets makes this a daunting task. Here we present Post-Acquisition Targeted Searches (PATS), a search tool that allows researchers without extensive proteomics knowledge and large computational resources to rapidly identify novel peptide and protein sequences in large published mass spectrometry data sets (>terabytes). With two of the largest interactome studies pre-loaded (5 terabytes of raw mass spectrometry data in total), PATS automatically visualizes protein correlation matrixes and protein network maps for all the bait proteins of each searched peptide sequence (e.g. Figure 2a-b), provides readily testable hypotheses for protein function based on the “guilt by association” concept”.

**Figure 1.**
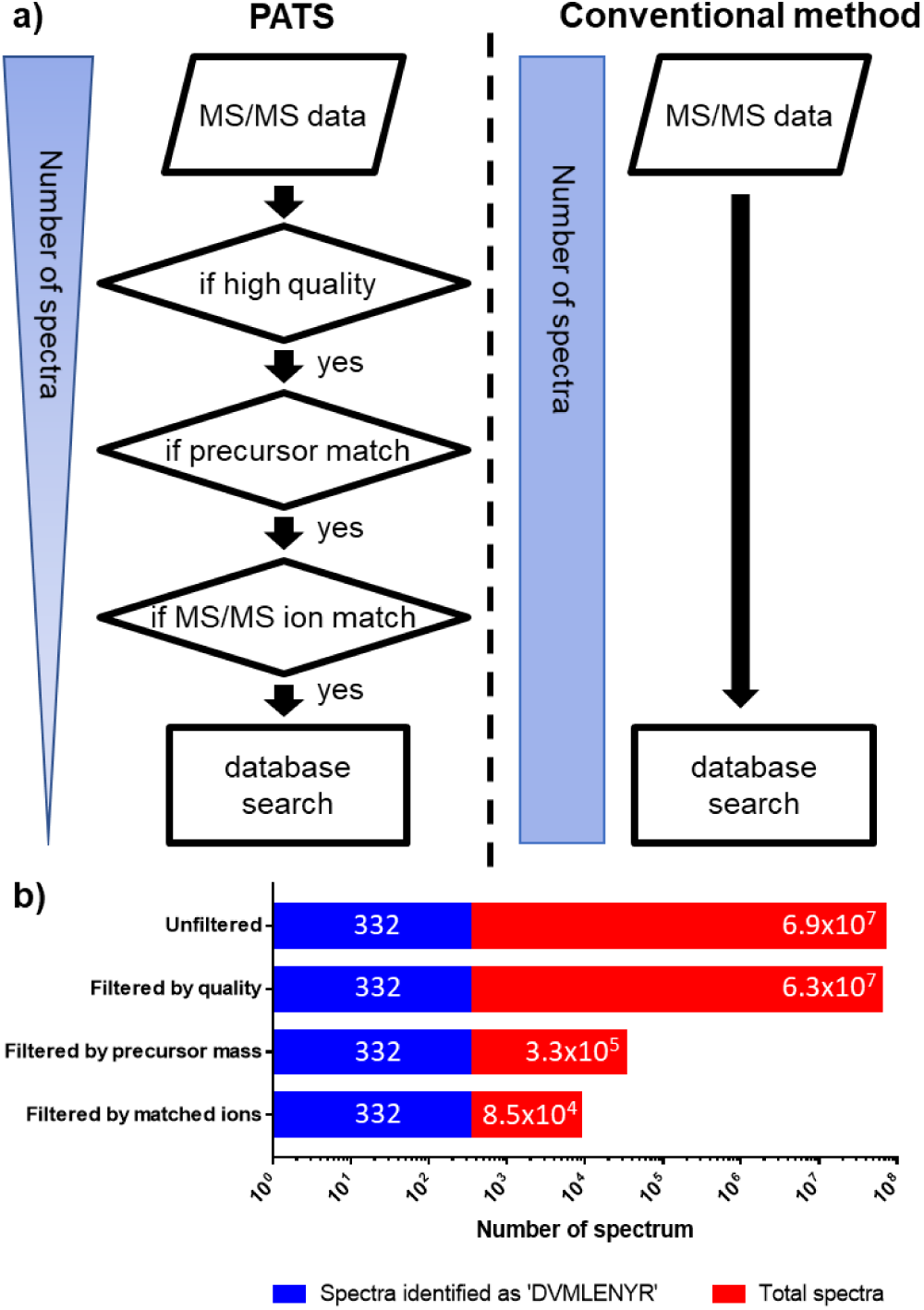
PATS data reduction by different stages. a) General scheme of the PATS method. b) Data reduction by PATS on peptide ‘DVMLENYR’ by different stages. PATS reduced number of spectra from 6.9×10^7^ to 8.5×10^4^.

The intent of the method is to provide an effective means to determine if a unique sequence (or other sequences), with or without PTMs, is present in large mass spectrometry data sets. In contrast to existing targeted strategies such as “peptide-centric proteome analysis” ^11, 12^ and “targeted database analysis” ^13^, which focused on analyzing either a single peptide or a focused group of peptides from all MS/MS spectra, PATS focuses on reducing the number of irrelevant MS/MS spectra being fed to database search algorithm (Figure 1b). Our strategy is especially optimized to work effectively with large data sets at terabytes scale. While all other existing strategies require at least hours to simply read terabytes of data from disk, PATS can extract all MS/MS spectra relevant to a target peptide from 5 terabytes of data within seconds and finish the whole analysis within minutes. PATS only reduces the amount of irrelevant MS/MS data and it does not alter the peptide identification or FDR estimation process. Therefore, each spectrum is still evaluated against all possible forward and reverse peptide candidates from a full-length reference database. The FDR is estimated the same way as conventional bottom-up proteomics methods.

PATS is available online at http://sequest.scripps.edu/PATS with two of the largest interactome mass spectrometry data sets pre-loaded (5 terabytes of raw mass spectrometry data in total). To search any specific peptide with or without PTM within these two data sets, user can simply submit their peptide sequence of interest to PATS and the result will be automatically generated (e.g. Figure 2a-b). For more advanced usage, the source code as well as a detailed instruction of how to run PATS on custom data set is available at https://github.com/bathyg/PATS/. To speed a search, PATS uses three methods for data reduction (Figure 1a, Supplementary Figure 1). In the first step, low-quality spectra that often result in unreliable peptide spectrum match (PSM) were eliminated. Any spectrum containing less than 20 peaks or with an average peak signal-to-noise ratio (SNR) less than 2.0 was excluded. In the second step, MS/MS data was filtered by the precursor ion m/z value of each spectrum. A precursor ion m/z value candidate list was generated for the target peptide, providing all possible charge states of the precursor ion. Spectra with precursor ion m/z values outside of the candidate list were excluded immediately and no longer processed. In the final step, comparing the theoretical fragment ions of the target peptide to each spectrum filtered all the remaining spectra. To do this a fragment ion list containing all *b* and *y* ions was calculated for each input peptide sequence and then at least five **b** or **y** ions have to match within 100 ppm of the fragment ion masses. After these three data reduction stages, the remaining tandem mass spectra are searched against the desired database.

**Figure 2.**
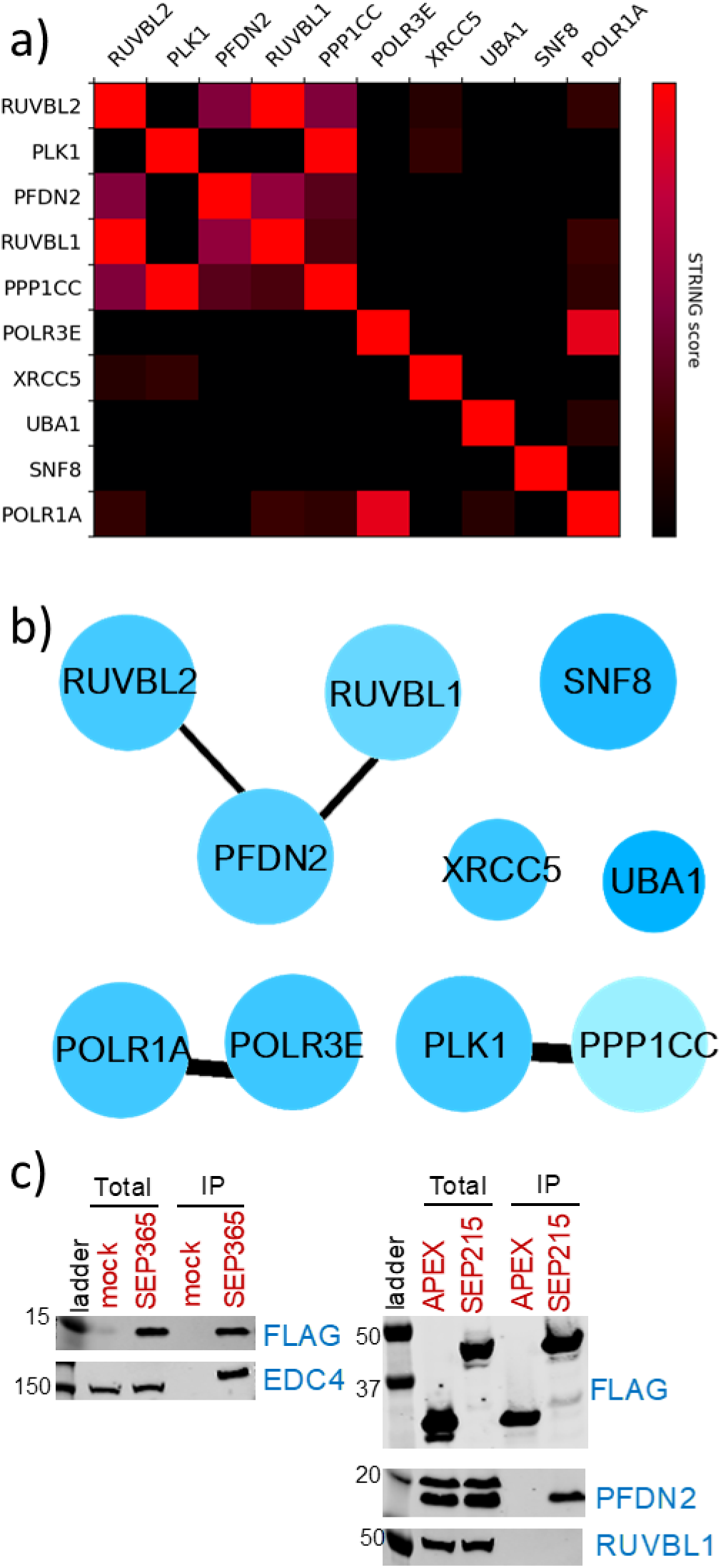
PATS detection and biochemical validation of SEP215. a) Protein correlation matrix generated by PATS for SEP215. b) Protein interaction network generated by PATS for SEP215. c) Western blot confirmation of SEP365 (positive control) and SEP215. Protein interactor of SEP215, PFDN2, was predicted by PATS and confirmed by our pull-down experiment.

To illustrate the process performed by PATS, a tryptic peptide sequence from the Krüppel associated box (KRAB) domain, ‘DVMLENYR’, was used as a peptide of interest and searched within a human interactome data set that consists of 4,293 AP-MS experiments containing more than 69 million spectra. When directly searched with a conventional search method using a reference protein database (Uniprot 2017-01, human, both reviewed and unreviewed), the search costs more than 40,000 CPU-hours (defined as total running hours × number of cores per CPU) to identify 19,521,327 peptide-spectrum matches (PSMs), of which 332 PSMs were related to peptide sequence ‘DVMLENYR’ (Table 1). Next, we used PATS with the same target peptide and database. The whole data set was filtered through PATS and then searched with equivalent parameters. The first step assesses the quality of tandem mass spectra and eliminates poor quality spectra. Of the 69 million spectra in the data set, 91% or 63 million pass the quality test (Figure 1b). The second step of filtering using precursor ion filter reduces the number of spectra to 32,904. Finally, in the third step, the number of spectra was further reduced to 8,452 spectra by matching predicted b and y ions. PATS required a total of 3 minutes 15 seconds to reduce 69 million spectra to 8,452. Then, when searched with a conventional algorithm against the same protein reference database with equivalent parameters, we identified 5,216 peptide-spectrum matches (PSMs) from PATS filtered data (8,452 spectra). From the 5,216 PSMs identified, we found the exact same 332 PSMs related to peptide sequence ‘DVMLENYR’ that was found with the conventional method. Overall, the search took less than 1 CPU-hour and obtained the same result as the conventional database search for peptide ‘DVMLENYR’, a speed increase of approximately 40,000-fold. This result showed that PATS filtered data were specifically enriched, from 0.0017% to 6.4% of total PSMs (3,765-fold enrichment), for our peptide of interest ‘DVMLENYR’ (Table 1) and therefore improved search efficiency.

**Table 1.**
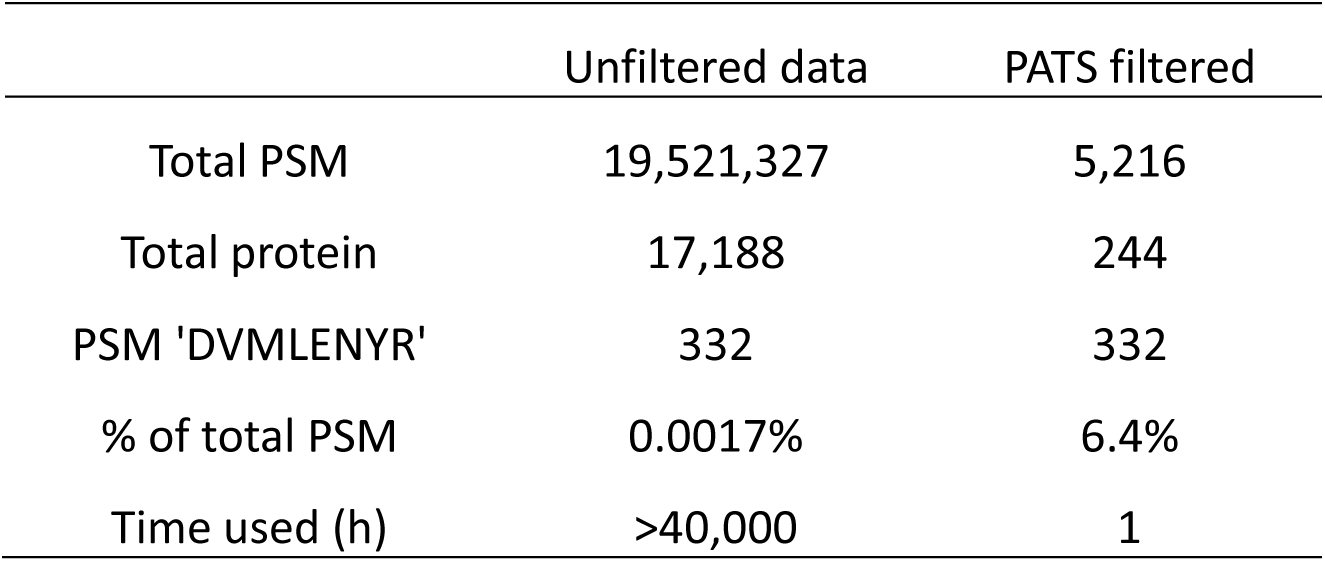
Number of identification of ‘DVMLENYR’ comparison. Number of identification of ‘DVMLENYR’ comparison peptide. PATS filtered data has significantly higher percentage of PSM (6.4% vs 0.0017%)

|  | Unfiltered data | PATS filtered |
| --- | --- | --- |
| Total PSM | 19,521,327 | 5,216 |
| Total protein | 17,188 | 244 |
| PSM 'DVMLENYR' | 332 | 332 |
| % of total PSM | 0.0017% | 6.4% |
| Time used (h) | >40,000 | 1 |

One application of PATS is to identify protein-protein interactions for proteins from previously non-annotated ORFs. In particular, the use of proteogenomics has uncovered the existence of hundreds to thousands of non-annotated microproteins derived from small ORFs (smORFs) in genomes from organisms ranging from yeast to humans. To date, the functions of almost of all of these proteins still remains a mystery. Determining if the products of these microproteins are functional is a difficult challenge and simply testing possible functions without a starting point is costly (e.g., large-scale screens). Instead, if some preliminary functional biochemical data could be obtained for these smORFs, it should be possible develop reasonable hypotheses that could speed their functional characterization.

A means to help assign putative functions to proteins is the “guilt by association” paradigm, which argues if a protein of unknown function interacts with a protein of known function then that protein’s function should be related^14^. The identification of microprotein-protein interactions has proven to be a successful approach for characterizing novel microproteins in a few cases. For example, the identification of protein-binding partners of a microprotein called NoBody revealed a role for this protein in modulating mRNA decapping^4^. To perform and search thousands of immunoprecipitation experiments to characterize microproteins is a daunting task and consequently PATS can be used to accelerate this process. To test the ability of the method to identify new biological information for small ORF encoded microproteins (SEP, sequence < 150 amino acids) in a large interactome, we searched 365 microproteins that had been previously identified by shotgun proteomics^15–17^. We first constructed a protein sequence database by appending the protein sequence of 365 microproteins (see Supplementary Table 1) to a Uniprot reference database (2017-01, human, both reviewed and unreviewed). Reversed sequences of all protein and common contaminants were then added automatically. This database was searched with MS/MS from the 4,293 AP-MS experiments^10^ using a conventional database search algorithm^18^ on a 200-node cluster. The search took in total 43,258 CPU hours to finish. In comparison, when PATS was used to filter the data prior to the search, the search finished within 224 CPU-hours (including PATS filtering time), or a speed increase of 200-fold compared to the conventional search. (Table 2). We then compared the results from both conventional search and PATS search. As shown in Table 2, there was virtually no difference in microprotein identifications, but significantly less for all other proteins, showing PATS successfully reduced search space without loss of confidence in the protein identifications. The percentage of microproteins at the protein and peptide level increased from 0.70% and 0.12% to 38.46% and 26.36% respectively, showing a significant enrichment. In total, we identified 120 microproteins (out of 365 microproteins) from this large data set with 231 peptides and 7642 PSMs (Table 2).

**Table 2.** Number of identification of SEP comparison. PATS significantly enriched SEP related identification on both protein (38.46% vs 0.70%) and peptide (26.36% vs 0.12%) levels.

|  | Normal search |  | PATs |  |
| --- | --- | --- | --- | --- |
| Total | Protein | Peptide | Protein | Peptide |
| Unique | 17188 | 187158 | 312 | 876 |
| SEP | 120 | 231 | 120 | 231 |
| PSM | 21093956 |  | 101366 |  |
| SEP PSM | 7655 |  | 7642 |  |
| SEP % | 0.70 | 0.12 | 38.46 | 26.36 |
| SEP PSM % | 0.04 |  | 7.54 |  |

In this human interactome MS/MS data set, each raw data file represents an AP-MS pull-down experiment from HeLa cell lysate by a specific bait protein^10^. Proteins identified from a specific AP-MS experiment represent all interactors of the bait protein, either specific or non-specific (e.g. noise). Within all the identified microproteins, many were identified from multiple AP-MS experiments of several bait proteins. We noted that many of the bait proteins belong to the same protein family or have close connections. Therefore, it would be interesting to see if these potential interactors of each microprotein interact with each other and form protein interaction network. PATS automatically generated protein correlation matrixes of the bait proteins using STRING-DB (string-db.org) correlation score (Figure 2a.) and K-mean clustering algorithm. From the correlation matrix and the confidence of peptide identification, PATS then generated an interactive protein network representing all the bait proteins that interact with the target protein/peptide. (Figure 2b.) The size of each node represents identification confidence and the color represents identification frequency (total spectral count) in the whole data set. The thickness of edges between nodes represents the STRING-DB correlation between two proteins. As shown in Figure 2b, there were three distinct protein clusters. The one with most highly confident proteins and interactions was the RUVBL1-PFDN2-RUBVL2 cluster. In order to further validate this interaction, we custom synthesized 18 synthetic peptides and compared their MS/MS with the ones identified by PATS. Most of the synthetic peptides showed almost identical fragmentation pattern with the ones identified by PATS (Supplementary Spectra 1-18), showing peptide level evidence of the existence of these detected microproteins.

With these data in hand, we went on and validated the results of the identification for some microprotein-protein interactions using immunoprecipitation against FLAG epitope. We transiently transfected HEK293T cells with FLAG-tagged SEP365 (see Supplementary Table 1 for microprotein name and sequence used in this study) or empty vector and APEX-FLAG-tagged SEP215 or APEX-FLAG only vector. The anti-FLAG immunoprecipitates from cell lysates were analyzed by Western blot to quantify proteins specifically enriched over a vector only or APEX-FLAG immunoprecipitation control. SEP365 was one of the top hits in PATS, also recently reported to be involved in mRNA decapping complex^4^. The Western blot showed a consistent result where EDC4, a member of mRNA decapping complex, was enriched in SEP365-FLAG immunoprecipitation (Figure 2c). PATS identified SEP215 to be associated with RUVBL1/2 and PFDN2 protein complex with high confidence. We performed immunoprecipitation and blotted against both PFDN2 and RUVBL1. We identified that PFDN2 as a potential direct interactor as shown by strong enrichment of PFDN2 in a Western blot, however we did not detect enrichment of RUVBL1 (Figure 2c). This may suggest that SEP215 has strong association with RUVBL1/2 in a protein complex but does not directly bind. RUVBL1/2 and PFDN2 protein complex was reported to function in RNA polymerase II assembly. PATS successfully identified SEP215 as an additional component of this protein complex and this allows us to form a targeted hypothesis and to further elucidate the function of novel microproteins. A list of all the identified SEP proteins and the corresponding peptides can be found both online at http://sequenst.scripps.edu/PATS and in Supplemental Table 2.

In summary, PATS provides an efficient method to search peptide or protein sequences of interest from large published proteomics data sets. Using interactome data sets, PATS visualizes the related protein interaction network and is helpful for the assignment of a putative function based on the “guilt by association” concept. By sequentially filtering MS/MS data by data quality, precursor mass, and MS/MS fragment ions, PATS significantly reduced the computational complexity and time required for database search of targeted peptides. We have shown that PATS can be used to search individual peptide as well as group of novel proteins with very high efficiency. The protein correlation matrix and protein interaction map generated by PATS can be used to identify novel protein interactors of protein/peptide of interest. Currently, PATS is preloaded with two of the largest human protein interaction data sets^8–10^ and can be used freely at http://sequenst.scripps.edu/PATS. We hope to include interactome data sets from other human cell lines as well as other species in the near future.

## ACKNOWLEDGMENTS

This work was performed at The Scripps Research Institute and Salk Institute for Biological Studies with the financial support from: The National Institutes of Health to J.R.Y (R01 MH067880, P41 GM103533, U54 GM114833), National Institutes of Health to A.S. (P30 CA014195 MASS core, R01 GM102491), Dr. Frederick Paulsen Chair/Ferring Pharmaceuticals to A.S. and Larry Hillblom Foundation to J.M. We thank the Mass Spectrometry Core Facility at Salk for providing access to their equipment and support.

## AUTHOR CONTRIBUTIONS

Y.G., J.M., A.S. and J.R.Y. designed the experiments. Y.G. developed the algorithms and implemented the codes and website. J.M. performed biochemical validations. Y.G., J.M., A.S. and J.R.Y wrote the manuscript.

## COMPETING FINANCIAL INTERESTS

The authors declare no competing financial interests.

## METHODS

### Raw data conversion

Raw data was obtained from Pride archive (https://www.ebi.ac.uk/pride/archive, PXD002815) and Bioplex website (http://bioplex.hms.harvard.edu). The raw files were first converted by rawconverter (http://fields.scripps.edu/rawconv/) to ms1 and ms2 files, then processed by PATS (https://github.com/bathyg/PATS) into indexed files.

### PATS usage

To use the pre-loaded PATS with either human interactome dataset^10^ or Bioplex interactome dataset^8, 9^, user should first go to the website (http://sequest.scripps.edu/PATS/) and follow the instruction. First, user can query PATS by gene name or tryptic peptide sequence without PTM (check partial peptide sequence checkbox to match partial sequence). If already searched and deposited in the database, PATS will return the protein network and protein interaction correlation matrix. If not found, user can then use the submit form to submit a custom search by specifying the tryptic peptide sequence and PTM. For example, “DKRPGSLETC (57.02146) R” will search peptide sequence DKRPGSLETCR with a specific PTM of 57.02146 Da on cysteine. Once completed, a link to the result will be send to the provided email address. For advanced usage, detailed documentation on PATS usage can be found at https://github.com/bathyg/PATS together with the source code.

### Cell culture and transfection

HEK293T cells were culture in DMEM supplemented with 10% fetal bovine serum. Cells were maintained in a 5% CO2 atmosphere at 37°C. Plasmid transfection was performed with Lipofectamine 2000 and Opti-MEM according to the manufacturer’s instructions. For co-immunoprecipitation, cells transfected with plasmids were assayed 24 hours post-transfection.

### Co-immunoprecipitation

FLAG-tagged SEP365 in pcDNA3 (or empty pcDNA3 vector as a negative control), and APEX-flag-tagged SEP215 in pcDNA3 (or APEX-flag-pcDNA3 as a negative control) were transfected into HEK293T cells using 10 µg DNA per 10 cm dish of cells. 24 hours post-transfection, cells were harvested and lysed using IP lysis buffer (Pierce) supplemented with Roche Complete protease inhibitor cocktail tablets. 1 mL lysis buffer was used per pellet. Cells were lysed on ice for 5 min followed by sonication in water bath for 5 min, then centrifugation at 17,000 g, 4 °C, 15 min. Lysate samples (20 ul) were saved for Western blot analysis. A 100 μL aliquot of anti-FLAG agarose beads (clone M2, Sigma) was washed with 1 mL of the lysis buffer, collected by centrifugation for 1 min at 3000 rpm, then suspended in the cell lysate supernatant. Bead suspensions were rotated at 4 °C for 2 hours, then washed 3 times with TBS-T. Elution was in 180 μL of 3X FLAG peptide (Sigma), at a final concentration of 125 μg/mL in TBS-T at 4 °C for 1 hour. Beads were removed by centrifugation and the supernatant was collected for Western blot analysis.

### Western blot

Cell lysates and eluents were loaded on a Bolt 4–12 *%* BisTris gel, 10-well (Life Technologies) and run in MES running buffer at 200V for 20 min. Proteins were transferred to PVDF membrane using iBLOT 2 (Life Technologies) program “P0”, followed by blocking the membrane at room temperature for 1 hour. Then the membrane was blotted with primary antibody: mouse anti-FLAG M2 (Sigma), rabbit anti-EDC4 (CST), rabbit anti-PFDN2 (Abcam) or rabbit anti-RUVBL1 (CST) at 1:1000 dilution at 4 oC overnight, Wash membrane three time with TBS-T, then blot with secondary antibody: goat anti-rabbit IRDye 800CW (LiCor) or goat anti-mouse IRDye 800CW (LiCor) at 1:10000 dilution, rock 1 hour at room temperature. Wash membrane three times with TBS-T then scan the membrane using LiCor Odyssey CLx at IR700 and IR800.

## REFERENCES

1. Eng, J.K., McCormack, A.L. & Yates, J.R. An approach to correlate tandem mass spectral data of peptides with amino acid sequences in a protein database. J Am Soc Mass Spectrom 5, 976–989 (1994).

2. Cheng, H. et al. Small open reading frames: current prediction techniques and future prospect. Curr Protein Pept Sci 12, 503–507 (2011).

3. Couso, J.P. & Patraquim, P. Classification and function of small open reading frames. Nat Rev Mol Cell Biol 18, 575–589 (2017).

4. D’Lima, N.G. et al. A human microprotein that interacts with the mRNA decapping complex. Nat Chem Biol 13, 174–180 (2017).

5. Saghatelian, A. & Couso, J.P. Discovery and characterization of smORF-encoded bioactive polypeptides. Nat Chem Biol 11, 909–916 (2015).

6. Kim, M.S. et al. A draft map of the human proteome. Nature 509, 575–581 (2014).

7. Wilhelm, M. et al. Mass-spectrometry-based draft of the human proteome. Nature 509, 582–587 (2014).

8. Huttlin, E.L. et al. Architecture of the human interactome defines protein communities and disease networks. Nature 545, 505–509 (2017).

9. Huttlin, E.L. et al. The BioPlex Network: A Systematic Exploration of the Human Interactome. Cell 162, 425–440 (2015).

10. Hein, M.Y. et al. A human interactome in three quantitative dimensions organized by stoichiometries and abundances. Cell 163, 712–723 (2015).

11. Gillet, L.C. et al. Targeted data extraction of the MS/MS spectra generated by data-independent acquisition: a new concept for consistent and accurate proteome analysis. Mol Cell Proteomics 11, O111 016717 (2012).

12. Ting, Y.S. et al. Peptide-Centric Proteome Analysis: An Alternative Strategy for the Analysis of Tandem Mass Spectrometry Data. Mol Cell Proteomics 14, 2301–2307 (2015).

13. Shanmugam, A.K. & Nesvizhskii, A.I. Effective Leveraging of Targeted Search Spaces for Improving Peptide Identification in Tandem Mass Spectrometry Based Proteomics. J Proteome Res 14, 5169–5178 (2015).

14. Hazbun, T.R. et al. Assigning function to yeast proteins by integration of technologies. Mol Cell 12, 1353–1365 (2003).

15. Ma, J. et al. Improved Identification and Analysis of Small Open Reading Frame Encoded Polypeptides. Anal Chem 88, 3967–3975 (2016).

16. Ma, J. et al. Discovery of human sORF-encoded polypeptides (SEPs) in cell lines and tissue. J Proteome Res 13, 1757–1765 (2014).

17. Slavoff, S.A. et al. Peptidomic discovery of short open reading frame-encoded peptides in human cells. Nat Chem Biol 9, 59–64 (2013).

18. Xu, T. et al. ProLuCID: An improved SEQUEST-like algorithm with enhanced sensitivity and specificity. J Proteomics 129, 16–24 (2015).

